# The hidden phase of memory: EEG signatures of reactivation during post-retrieval rest

**DOI:** 10.1101/2025.11.03.685924

**Authors:** Luz Bavassi, Germán Campos-Arteaga, Ismael Palacios-García, Mario Villena-Gonzalez, Libertad Campassi, Emiliano Marachlian, Cecilia Forcato, Eugenio Rodriguez Balboa, Eugenia Pedreira

**Affiliations:** Instituto de Fisiología, Biología Molecular y Neurociencias (IFIBYNE), CONICET, Buenos Aires, Argentina; Departamento de Física, Universidad de Buenos Aires, Buenos Aires, Argentina; Departamento de Ciencias Jurídicas y Sociales, Facultad de Ciencias Jurídicas y Sociales, Universidad Tecnológica Metropolitana, Santiago, Chile; Centro de Estudios en Neurociencia Humana y Neuropsicología, Facultad de Psicología, Universidad Diego Portales, Santiago, Chile; Laboratorio de Sueño y Memoria, Instituto Tecnológico de Buenos Aires (ITBA), Buenos Aires, Argentina; Pontificia. Universidad Católica de Chile, Laboratorio de Neurodinámica Básica y Aplicada, Escuela de Psicología, Santiago, Chile

**Keywords:** Resting State, Memory reactivation, Beta power reduction, Subsequent memory accuracy, Phase synchronization network

## Abstract

Memory retrieval reactivates previously encoded representations, allowing for their modification and strengthening. However, the neural processes that follow reactivation and contribute to long-term retention remain poorly understood. Here, we examined the post-retrieval resting period to identify neural markers of memory reactivation and their relation to subsequent memory performance. Participants learned pairs of nonsense syllables in multisensory contexts and, on the following day, received either a cue-syllable reminder (context + cue syllable; RX) or a context-only reminder (RCTX) while EEG activity was recorded before and after the reminder presentation. Both reminders elicited significant reductions in beta power (25–40 Hz) during the post-reminder rest, consistent with memory reactivation. The magnitude of beta decrement correlated with better long-term performance. Graph-theoretical analyses of phase synchronization networks in the beta band revealed that the RCTX reminder produced higher bilateral frontal betweenness centrality, suggesting greater engagement of frontal regions in mediating global information flow when retrieving only contextual cues. Moreover, frontal centrality and network density were predictive of subsequent memory accuracy. These findings demonstrate that memory reactivation extends beyond cue presentation into post-retrieval rest, leaving identifiable oscillatory and topological signatures that influence memory persistence. Our results underscore the crucial role of frontal network dynamics and beta-band activity in facilitating long-term memory.

## 1 Introduction

Memory updating is a crucial aspect of adaptability in both humans and non-human animals, enabling flexible behavior in a constantly changing environment. The presentation of external cues triggers memory retrieval, re-expressing specific aspects of the spatiotemporal neural activity pattern that occurred during the initial experience Josselyn and Frankland [2018]. This reactivation has been proposed to mediate memory malleability by opening a time window during which the original memory trace can be modified Frankland and Bontempi [2005]. Depending on the information embedded in the cue and the characteristics of the original memory, its content can be updated, strengthened Ye et al. [2020], or weakened Wimber et al. [2015]. Although memory malleability is fundamental to everyday life, the underlying neural mechanisms remain largely unexplored.

Memory reactivation is not confined to a single moment; different neural signatures contribute across multiple timescales during retrieval. Within the first seconds after cue onset, the level of neural reactivation predicts both immediate Johnson et al. [2015] and long-term recall performance Kang et al. [2022]. Considering that neural reinstatement also occurs during rest periods after encoding—facilitating memory retention Tambini and Davachi [2019]—it is reasonable to assume that this mechanism extends beyond the retrieval period. Supporting this notion, several studies suggest that spontaneous activity during post-retrieval rest reflects memory updating Feng et al. [2015] and memory generalization Liu et al. [2019]. Conversely, presenting new information immediately after a reminder increases long-term errors compared to presenting it before the reminder Sinclair and Barense [2018]. These findings indicate that brain activity during rest periods following cue presentation may critically influence long-term memory performance.

Brain oscillations play a central role in long-term memory processes, including consolidation, reactivation, and retrieval Fell and Axmacher [2011], Schreiner and Staudigl [2020]. In episodic memory, decreases in low-frequency power — particularly in the alpha/beta range (8 − 30Hz) — have been associated with successful encoding and retrieval, reflecting reduced neural noise and enhanced information processing across sensory modalities Hanslmayr et al. [2016, 2019], Griffiths et al. [2019], Waldhauser et al. [2016], Hanslmayr et al. [2011]. While power decreases are not stimulus-specific, oscillatory phase can convey task- and content-related information: during sensory reinstatement, alpha/beta phase patterns encode stimulus-specific details, often accompanied by concurrent power reductions in the same regions Ng et al. [2013], Michelmann et al. [2016]. Thus, the power and phase of alpha/beta oscillations provide complementary neural signatures of successful memory retrieval.

Like other higher cognitive processes, memory relies on the anatomical and functional interactions among distributed brain regions McIntosh [1999]. Neuroimaging studies suggest that memory depends on transient interactions between specific regions Cabeza and Moscovitch [2013]. Recent findings indicate that presenting a fear reminder strengthens the functional connectivity between the amygdala and ventromedial prefrontal cortex during the subsequent resting period, highlighting its relevance for memory updating Feng et al. [2016]. Moreover, we previously showed that reactivating an episodic memory leaves distinct global network signatures during long-term retrieval, depending on the type of reminder presented Bavassi et al. [2019]. Altogether, these results underscore the need to further investigate the role of functional connectivity following reminder presentation and its influence on changes in stored information.

The main goal of the present study is to characterize the neural signatures of memory reactivation during a post-reminder resting period and to determine their relationship with long-term behavioral performance. To this end, we employed a well-established behavioral paradigm developed in our laboratory. The task follows a three-day protocol in which participants learn five pairs of nonsense syllables, each presented within an enriched multisensory context (image and music) Forcato et al. [2007, 2009, 2011], Rodríguez et al. [2013]. Using electroencephalography (EEG), we analyzed resting periods following two types of reminders presented on Day 2: one that included the contextual information plus the cue syllable (RX reminder) and another that included only the contextual information (RCTX reminder). Specifically, we assessed the frequency power and phase-synchronization functional connectivity induced by each reminder and examined their impact on long-term memory performance.

## 2 Results

To characterize the neural signatures of memory reactivation associated with long-term memory performance, we implemented a syllable-learning protocol comprising three separate sessions Forcato et al. [2007, 2009, 2011], Rodríguez et al. [2013].. During the Training Session, participants learned five pairs of nonsense syllables presented in a rich multisensory environment that included music and images. They completed ten trials to encode the syllable pairs. The following day, during the Treatment Session, participants were reminded of the information learned on the first day. Finally, their memory retention was assessed during the Testing Session (Figure 1A). On Day 2, we recorded EEG activity pre- and post-presentation of the reminder (*bl*1 and *bl*2, respectively). We recorded two resting-state periods: a 4-minute baseline rest before the reminder and a 4-minute post-reminder rest to examine any lingering neural effects. Participants were assigned to one of two experimental groups. The first group received the RX reminder, which included contextual information and the cue syllable, and was interrupted before participants could write the response syllable. In contrast, the other group received the RCTX reminder, which consisted solely of the contextual cues from the learning environment.

**Figure 1:**
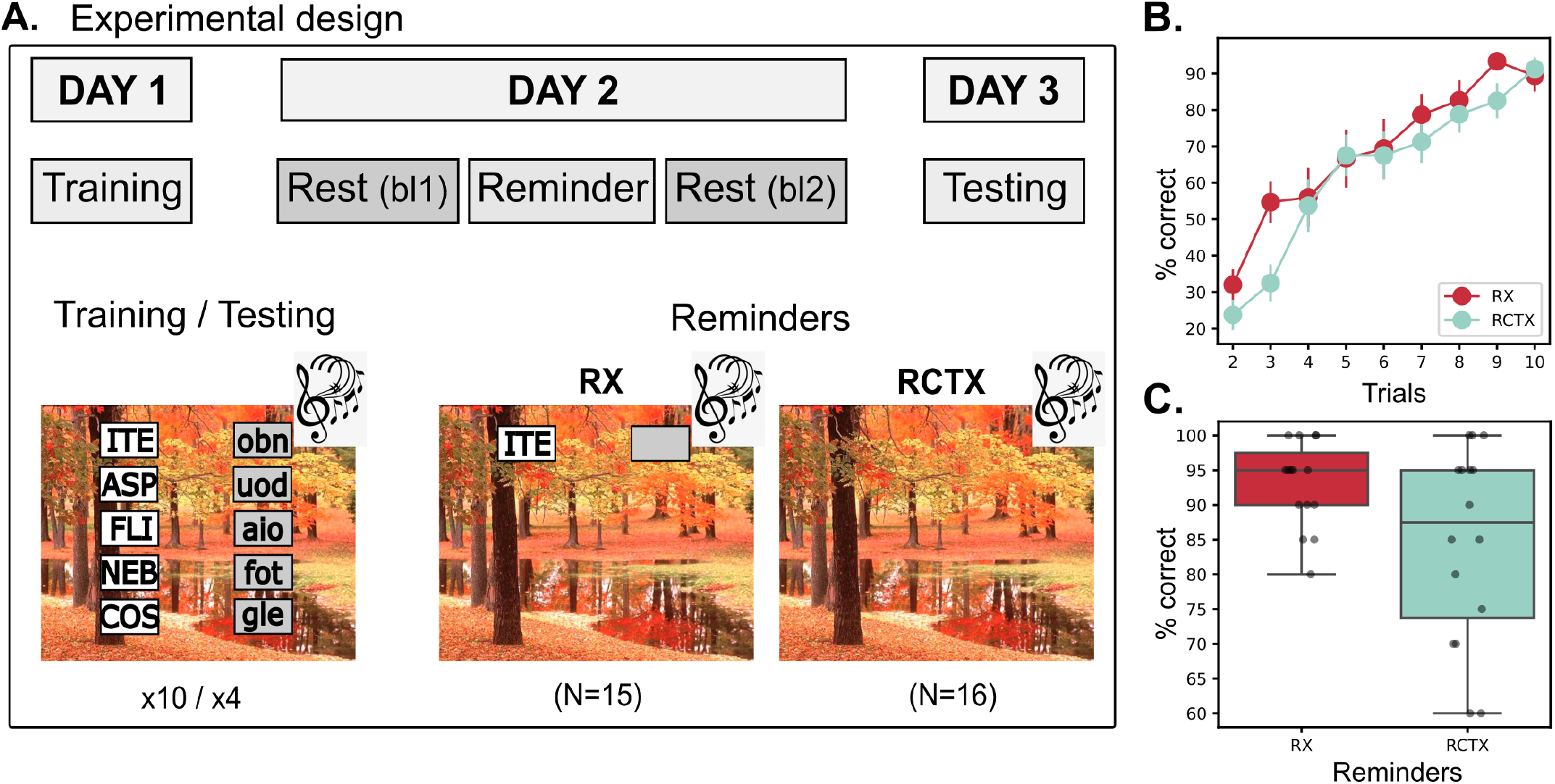
Overview of the syllable paradigm and behavioral results. **A**. The experimental paradigm follows a three-day task in which participants learn five pairs of nonsense syllables, each presented within an enriched, multisensory context (comprising an image and music). Each pair consists of a cue syllable associated with a response syllable. On Day 1, participants learnt the syllables by repeating each trial 10 times. On day 2, each participant was randomly assigned to an experimental group. The first one received the RX reminder (N=15), which consisted of the context plus one cue syllable. The reminder presentation finished before the participant responded. The other group received the RCTX reminder (N=16), which contained only contextual information. Before (*bl*_1_) and after (*bl*_2_) the reminder presentation, we recorded 5 minutes of resting-state to assess neural activity. Finally, on Day 3, we evaluated long-term performance by replicating 4 Training Session trials. **B**. The learning curve for each group on Day 1 **C**. The percentage of correct responses on the Testing Session on Day 3. (RX: red; RCTX: green).

First, we characterized the behavioral results of the task. The learning rate was similar across groups; participants improved their performance across training trials, reaching near-perfect scores in the final two trials (Figure 1B). During the Testing Session, we observed a high memory retention for both groups. The long-term memory test did not show significant differences between the mean percentage of correct responses for both groups (*U* = 158, *p* = 0.06, Mann–Whitney test, *effect size* = 0.7; Figure 1C). Notably, the two groups displayed different variances in their performance. Levene’s test confirmed a significant difference in variance between the groups (*F* (2, 29) = 9.02, *p* = 0.006), with the RX reminder group showing substantially lower variability. These results indicate that both reminders strengthen memory, counteracting forgetting; however, the reminder that includes the syllable and context (RX) appears to have a greater effect on long-term retention, as reflected in reduced variability.

This experiment aimed to identify neural signatures of memory reactivation, particularly those associated with memory enhancement. To this end, we first examined the neural markers elicited by reminder presentations in each experimental group. We applied a cluster-based permutation test across spatial and spectral dimensions to compare resting-state EEG activity after (*bl*_2_) and before (*bl*_1_) each reminder presentation (Figures 2A and 2B). Surprisingly, the overall spectral patterns observed across the two experimental groups were quite similar, suggesting that both types of reminders produced comparable frequency-power signatures. In both groups, the reminders triggered a significant decrease in beta power. A closer inspection of the significant clusters revealed differences in their topographical distribution. The group that received the RX reminder exhibited a localized cluster in the left parietal–occipital region between 25 and 45Hz (*p* = 0.05, *t* = − 353.5,cluster-based permutation test). On the other hand, the group receiving the RCTX reminder showed a bilateral central cluster, including frequencies from 20 Hz to 45 Hz (*p* = 0.02, *t* = − 565.7, cluster-based permutation test).

**Figure 2:**
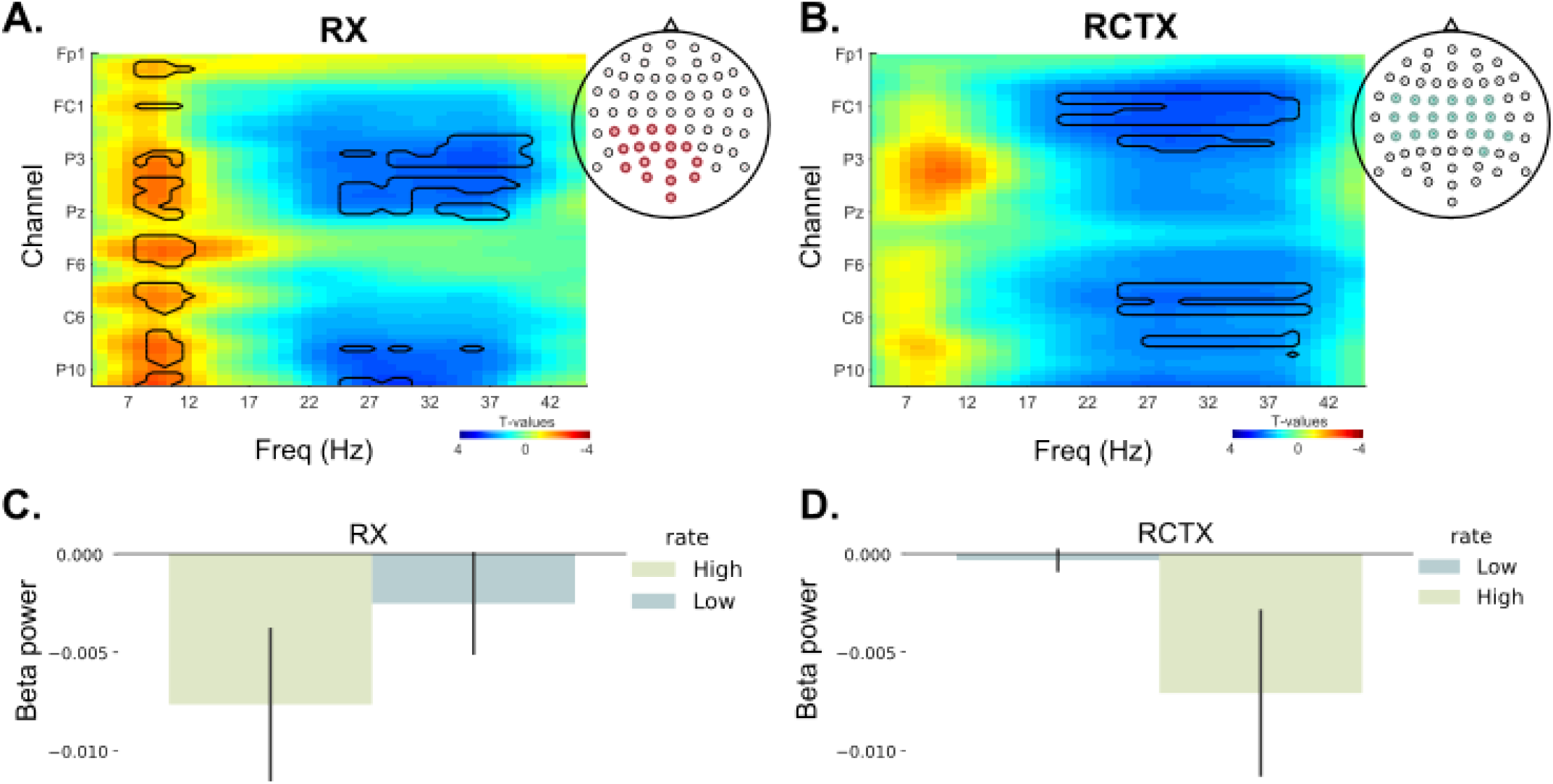
Spectral power analysis. **A**. T-maps of spectral power changes between *bl*_2_ and *bl*_1_ in the RX group. The thick black line depicts the significant cluster (*p <* .05, cluster-based permutation test). On the left, the location of EEG-channels for the beta cluster. **B**. T-maps of spectral power changes between *bl*_2_ and *bl*_1_ in the RCTX group. the thick black line depicts the significant cluster (*p <* .05, cluster-based permutation test). On the left, the location of EEG-channels for the beta cluster. **C** and **D**. The *bl*_2_ − *bl*_1_ power of the significant cluster for High and Low performance in the Testing Session, for the RX reminder group and the RCTX reminder group, respectively. (Low: blue; High: light green)

To further explore the relationship between beta power reduction and long-term memory performance, we divided each group according to the percentage of correct responses in the testing session (low vs. high performers, Figure SF1). We evaluated the magnitude of beta power reduction of each significant cluster (Figures 2C and 2D). In both groups, participants with higher testing performance exhibited greater decreases in beta power. This effect was only significant in the RX reminder group (RX: *U* = 40, *p* = 0.02; RCTX: *U* = 47, *p* = 0.10 Mann–Whitney test), probably indicating a higher effect of this reminder, although the sample size per condition is too small to ensure this result. In summary, both types of reminder presentations elicited reductions in beta power, linked to subsequent successful long-term memory performance. The RX reminder produced a parietal beta power decrement, while the RCTX reminder induced a beta power reduction over central regions. Since alpha/beta power reductions are considered proxies for information processing and are associated with successful subsequent retrieval, we interpret this to mean that both reminders trigger memory processing. Still, based on the topological features of the clusters, the two reminders engage different aspects of the previously learned declarative memory: the RX reminder may be reactivating more episodic information, while the RCTX reminder may be engaging semantic content.

Not expected, the RX reminder also revealed a widespread, significant cluster associated with an increase between (7 − 12*Hz, p* = 0.05, *t* = 293.8, cluster-based permutation test, Figures 2A). To understand this finding in terms of behavior, we again split this group’s participants by testing session (low vs. high performance) and did not observe a difference in alpha power (SF2; *U* = 24, *p* = 0.85; Mann–Whitney test). One possible interpretation of this result is that the presentation of the RX reminder is more effective in inducing participants to focus their attention. Interestingly, we performed a cluster-based permutation test comparing the post-reminder resting periods of both groups (*bl*_2_), and we did not find significant differences.

Our previous results suggest that both reminders—the RX and the RCTX—triggered memory reactivation and memory strengthening. To identify differences between the experimental groups, and considering that oscillatory phase conveys complementary information about memory and that functional connectivity plays a central role in memory processes, we examined the global network structure based on phase synchronization within the frequency range identified in the power analysis underlying each reminder. Specifically, EEG signals were bandpass filtered between 25 and 40 Hz. We then applied the Hilbert transform and computed the Phase Lag Index (*PLI*) between all pairs of channels Stam et al. [2007], generating a 64 *×* 64 symmetric matrix for each participant (see 4 for more details, Figure 3A).

**Figure 3:**
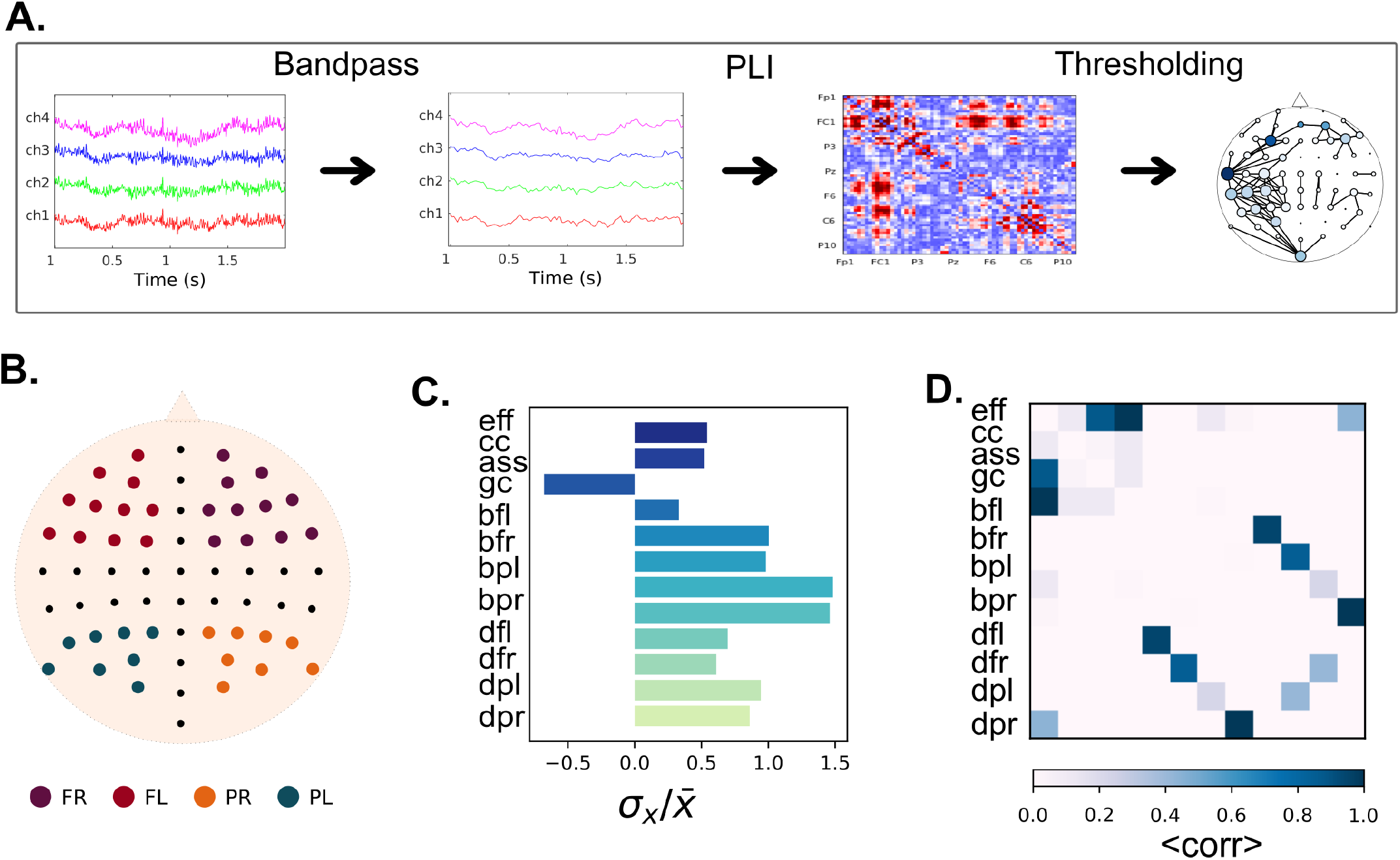
Phase-synchronization networks. **A**. Pipeline for building the brain’s network. First, we filtered the channel’s time series between 25 and 49Hz. Then, we computed the Phase Lag Index (PLI) for all channel pairs, yielding a 64 *times*64 symmetric matrix per subject. Finally, we thresholded each matrix, obtaining a binary adjacency matrix (i.e., a non-directed, unweighted graph) with densities ranging from 1% to 10%. **B**. We computed the degree and betweenness centrality for four regions: frontal right (FR), frontal left (FL), parietal right (PR), and parietal left (PL). **C**.The coefficient of variation (the standard deviation over the mean value) per each network measure (eff: efficiency; cc: clustering coefficient; ass: assortativity; gc: giant component; bfl: betweenness centrality of frontal left region; bfr: betweenness centrality of frontal right region; bpl: Betweenness centrality of parietal left region; bpr: Betweenness centrality of parietal right region; dfl: degree centrality of frontal left region; dfr: degree centrality of frontal right region; dpl: degree centrality of parietal left region; dpr: degree centrality of parietal right region) **D**. The correlation matrix between the network measures. Dark blue reflects a higher correlation coefficient.

To compare networks with the same density, we constructed binary, bidirectional graphs by thresholding each matrix, fixing the number of edges (Figure 3A). Densities ranged from *ρ* = 1% to *ρ* = 10%, yielding a set of 364 graphs per participant. At first glance, there was no possible way to distinguish networks for each condition (See Figure SF3). So, for each network, we computed four global attributes Barabási [2013]: Efficiency, which reflects the capacity for information exchange across the entire network; Assortativity, which measures the correlation in connectivity between nodes (whether nodes tend to connect to others with similar degree); Giant component size, which quantifies the number of connected nodes; and Clustering coefficient, indicating the extent to which a node’s neighbors are also interconnected. Additionally, we examined two nodal properties: Degree centrality, defined as the number of direct connections a node has; and Betweenness centrality, which captures the extent to which a node lies on the shortest paths between other nodes, thus reflecting its role in mediating information flow. Because we had no prior hypothesis regarding regional involvement, we divided the brain into four sections: bilateral frontal and parietal-occipital regions (Figure 3B).

In total, we computed 12 attributes per network, resulting in 4368 values per subject (12*x*364). To focus only on the informative features, we assessed the variability of each attribute across all networks, defined as the coefficient of variation (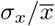; Figure3C). This analysis revealed that bilateral frontal betweenness centrality exhibited high variability and appeared to be a promising marker for distinguishing between experimental conditions. Next, recognizing that many graph-theoretical attributes are inherently correlated, we assessed pairwise correlations between features (Figure 3D). Efficiency showed strong correlations with both assortativity and giant component size, while betweenness centrality was highly correlated with degree centrality within each brain region. Based on these results, we selected six attributes for further analyzes: efficiency, clustering coefficient, and betweenness centrality (computed for the four brain regions).

Next, based on the set of six selected attributes, we examined the importance of these network measures in differentiating the two experimental conditions using a classifier model Breiman [2001]. The classification was performed across network densities ranging from 1% to 10%. Due to the limited data, we implemented a Random Forest classifier with 50 shallow trees (maximum depth of 2), each considering only two features at each split, to reduce the risk of overfitting. For each model, we developed a leave-one-out validation. Figure 4A presents the model’s mean accuracy across all network densities (the model’s accuracy on the training set was ∼ 0.85). We found that the classifiers performed above chance in 119 of the 364 network densities (33% of the times), especially for networks with densities of 4,6, and 8% (gray areas in Figure **??**B).

**Figure 4:**
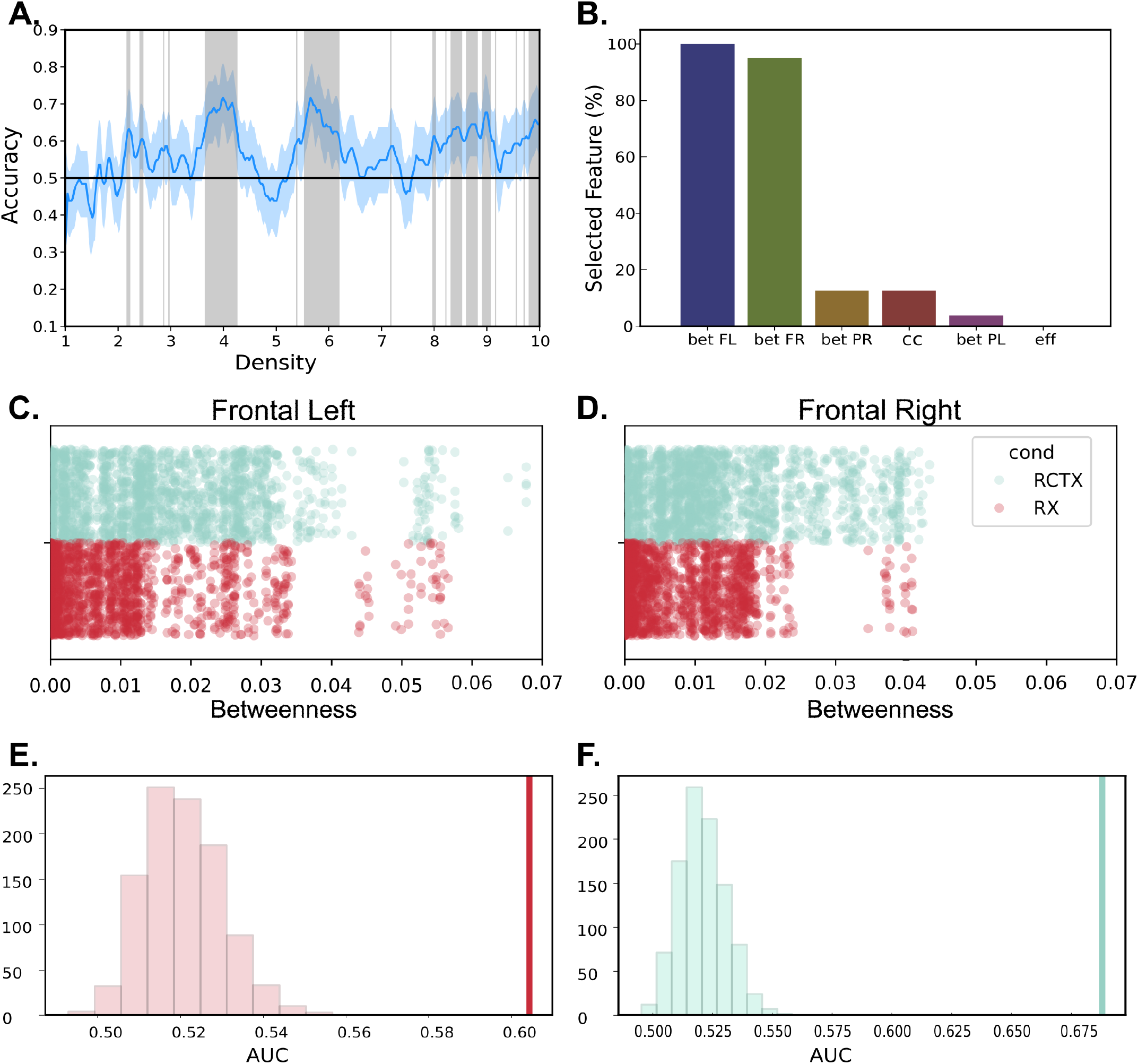
Feature importance approach. **A**. Accuracy of the Random Forest classifier in distinguishing the networks of both experimental groups for all network densities. Grey regions highlight the densities where the accuracy is above chance. **B**. The importance of the six features, sorted by decreasing importance. The left frontal betweenness centrality was the most important, followed by the right frontal betweenness centrality. Frontal left and right betweenness centrality for every network where the classifier was significant, **C** and **D**, respectively. **E**. Area under the ROC curve (AUC) for Logistic Linear Model to identify Low vs High performance in the RX reminder group by the contribution of left and right prefrontal betweenness and density. The line represents the AUC for experimental data, while the histogram illustrates the AUC for randomized data under the null model. **F**. AUC for the Logistic Linear Model in the RCTX (line) and the AUC for the model of the randomized data (histogram).

We analyzed the relative importance of six attributes in distinguishing between the two experimental conditions. For the graph densities where the classifier performed significantly above chance, we conducted model-based feature selection by computing permutation feature importance with 50 random shuffles. This method assesses how much the model’s performance drops when each feature is randomly permuted. Our analysis revealed that the frontal betweenness measures—both right and left—emerged as the most essential attributes for distinguishing the RX and RCTX networks.

Notably, the left frontal betweenness consistently ranked as the top feature across all densities. Additionally, the right frontal centrality was significant in 95% of the well-classified densities (Figure 4B).

Then, we visualized the centrality measure in both frontal regions for each subject and across all significant densities (Figures 4C and 4D). Interestingly, the networks associated with the RCTX reminder exhibited higher bilateral frontal betweenness centrality than those related to the RX (in both hemispheres; Mann-Whitney test: *p <* 0.001). Moreover, in both experimental groups, the left frontal region had higher betweenness centrality than the right one (RX: Mann-Whitney test: *p* = 0.0031; RCTX: Mann-Whitney test: *p <* 0.001). Finally, we performed a generalized linear model (GLM) to assess behavioral performance in the testing session (Low vs High performers) using both frontal betweenness centralities and the network link density per reminder. For the RX reminder, the overall model was statistically significant (Pseudo *R*^2^ = 0.099; *log*-*likelihood* = −1108.4). Among the predictors, frontal left centrality showed a strong positive effect on the probability of the outcome (*β* = 81.76, *z* = 10.15, *p <* 0.001), while density exhibited a significant negative association (*β* = −9.36, *z* = − 4.26, *p <* 0.001). In contrast, the frontal right centrality was not a significant predictor (*β* = − 4.16, *p* = 0.62). The intercept was also significant (*β* = 0.34, *p* = 0.006). To evaluate the model’s discriminative capacity, we calculated the area under the Receiver Operating Characteristic curve (AUC-ROC). Although it was moderate (AUC-ROC = 0.60), it was above-chance (Figure 4E). Besides, the linear model for the RCTX reminder provided a good overall fit (Pseudo *R*^2^ = 0.095; *log*-*likelihood* = − 1224.9). Both frontal centralities showed significant negative effects on the probability of the outcome (*β* = − 68.72, *z* = −12.31, *p <* 0.001; and *β* = −13.25, *z* = − 3.36, *p* = 0.001, right and left, respectively). In contrast, density had a significant positive effect (*β* = 14.24, *z* = 6.12, *p <* 0.001). The intercept was also statistically significant (*β* = 0.28, *p* = 0.018). Figure 4F displayed the AUC-ROC, which is above-chance, indicating reliable predictions in distinguishing the testing performance. Overall, bilateral frontal betweenness centrality is the primary feature that distinguishes the essence of the reminder. Considering that betweenness centrality reflects a node’s role in mediating information flow within a network, after the RCTX reminder, the frontal regions act as hubs, orchestrating global brain activity. Moreover, based on frontal network centrality and connectivity density, it was possible to predict subsequent performance in the Testing Session.

## 3 Discussion

In this work, we explored the neural signatures of reactivation in a resting period after the presentation of different reminders. We found that both reminders elicited a reduction in beta power (25 − 40 Hz), and this reduction was more pronounced among participants with low long-term performance in the Testing Session. Moreover, we found that the phase-synchronization networks induced by the RCXT reminder reflect a more prominent role for the frontal region, as it is more involved in mediating information flow across the network. Also, we showed that bilateral frontal centrality was sufficient to differentiate between the two reminder conditions.

Many memory paradigms focus on reactivation during encoding and retrieval. In this work, we propose a novel approach, investigating the post-retrieval rest period. In encoding, the processes unfolding during awake periods contribute positively to memory performance. The interruption of the endogenous reactivation during the post-encoding resting periods impaired memory retention Tambini and D’Esposito [2020]. Additionally, during this offline period, reactivation of successfully recalled objects is greater than that of forgotten ones Staresina et al. [2013]. The spontaneous activity post-retrieval was also related to the subsequent outcome, linked to fear memory updating Feng et al. [2015] and episodic memory generalization Liu et al. [2019]. Interestingly, our paradigm reveals that beta (25 − 40 Hz) power and phase act as neural signatures of memory reactivation during the resting period after both reminder presentations, indicating that reactivation extended beyond stimulus presentation. While power decrement is related to memory accuracy during the long-term testing session, the beta phase in frontal regions carries information about previously presented cues.

Hanslmayr and colleagues Hanslmayr et al. [2012] proposed that the reductions in alpha/beta power serve as a marker of memory reactivation. The authors argue that decreases in alpha/beta power serve as a proxy for enhanced neuronal firing, thereby facilitating the brain’s ability to represent information, reducing noise Hanslmayr et al. [2016]. In this sense, we observed that presenting both reminders led to a significant decrease in beta power (25–40 Hz), indicating that presenting either reminder induces memory reactivation. Moreover, a greater reduction in beta power was associated with better long-term accuracy for both reminders. This phenomenon has been reported during encoding, where decreases in alpha and beta power correlate with the number of items successfully retrieved Hanslmayr et al. [2009].

Interestingly, the reminders triggered clusters of significant decreases in beta power, which were differently localized across the brain. The RX reminder, which received the syllable plus the context information, presented a left parietal-occipital cluster, while the RCTX reminder (the one that only includes the context information) exhibited a bilateral central cluster. Recent evidence suggests that beta-band power reductions during memory retrieval are not uniform but instead reflect the modality and content of the memory. Griffiths et al. (2019) Griffiths et al. [2019] demonstrated that alpha/beta power decreases during successful retrieval of visual and auditory information are localized in modality-specific sensory areas: occipital cortex for visual memories and parietal/temporal cortices for auditory memories. These results suggest that memory retrieval involves reinstatement of activity in regions initially engaged during encoding. Supporting this view, a previous MEG study using crossmodal associative memory paradigms found that retrieval of an associated item (e.g., retrieving a visual image from an auditory cue) activated the sensory-specific cortex corresponding to the retrieved—not the presented—stimulus Wheeler et al. [2000], Waldhauser et al. [2016]. These findings converge to suggest that beta desynchronization serves as a signature of cortical reinstatement, reflecting the fidelity of retrieved sensory content. Our task involved multisensory information, comprising semantic, visual, and auditory information. In this sense, we proposed that the two reminders trigger distinct sensory aspects of the encoded memory. The RX group may primarily be reactivating visual information comprising semantic content due to left lateralization. In contrast, the RCTX reminder induces a power reduction in auditory cortex regions.

In contrast to previous studies, which propose that decreases in alpha power during the perception of sequential stimuli correlate with improved memory performance Griffiths et al. [2019, 2021], in this report, the RX group displayed a significant widespread cluster linked to an increase in alpha power. It is essential to note that the RCTX reminder group also showed, although not statistically significant, an increase in alpha power. Alpha activity was initially interpreted as a sign of cortical inactivity Berger [1929]. However, later studies demonstrated that alpha oscillations actively contribute to attention by inhibiting processing in task-irrelevant areas, thereby enhancing selective attention Kelly et al. [2006]. In line with these findings, as our paradigm did not explicitly direct participants to focus on any step of the protocol, we speculate that the RX group is focusing on completing the response syllable following the presentation of the cue syllable. This expectation is lower in the RCTX group because there is no such information.

Although both reminders prompt memory reactivation, as indicated by a decrease in beta power, we found subtle differences in behavioral performance. Considering that beta’s phase carries stimulus-specific memory, we explored global network phase synchronization, specifically aiming to distinguish between the two types of retrieval. Our analysis revealed that the frontal betweenness centrality is a key attribute in differentiating the two conditions. The RCTX reminder group presented higher frontal betweenness centrality than the RX reminder group. In other words, the RCTX reminder induces a network in which the left and right frontal regions play pivotal roles, serving as central stations for information transfer throughout the entire brain. This reminder provides fewer retrieval cues directly linked to the original learning than the RX, leading to diminished direct access to the original memory trace. As a result, retrieval in the RCTX group might entail increased internal searching, thereby requiring greater engagement of prefrontal cortex mechanisms. Neuroimaging studies have indeed indicated that activation of the dorsolateral prefrontal cortex (DLPFC) increases when memory strength is lower or when cues are insufficient, reflecting the need for controlled retrieval and attentional support Donaldson et al. [2010], Kirwan et al. [2008]. Furthermore, Kuhl et al. (2011) demonstrated that memory strength influences prefrontal cortex activation during retrieval, emphasizing that a reduced overlap between cues and memories correlates with greater retrieval effort Kuhl et al. [2011].

At this stage, we can interpret our findings within the framework of Nonmonotonic Plasticity Ritvo et al. [2019], which offers a compelling explanation for how varying degrees of memory reactivation can lead to distinct outcomes, including integration, no change, or differentiation of memory traces. This model provides a valuable perspective on the differential effects observed following the presentation of cue and context reminders in our study. We assume that memory retrieval triggers reactivation of the original memory trace through a diffusive propagation of activation Bavassi and Fuentemilla [2024]. Both types of reminders initiated memory retrieval by reactivating the original memory trace; however, they did so to varying extents. Notably, although the RX reminder is more closely aligned with the original learning episode than the RCTX reminder, this similarity alone is insufficient to account for the more substantial behavioral and neural effects observed. A critical aspect of our paradigm is that the RX reminder implicitly encourages participants to complete the learned association upon reading the cue syllable. This internal retrieval operation is likely to lead to greater reactivation. On the other hand, the RCTX reminder does not prompt this demand, leading to weaker reactivation. According to the nonmonotonic model, this difference in activation level may elucidate why the RX reminder promotes memory strengthening (through integration), while the context reminder yields weaker or even neutral behavioral outcomes Forcato et al. [2014]. Thus, our results indicate that, beyond stimulus similarity, task demands and the cognitive operations they evoke play a pivotal role in shaping reactivation strength and, ultimately, memory modification.

All in all, our study provides insight into the early neural dynamics underlying different types of memory reactivation during the resting state. This approach contrasts with previous studies, which have primarily focused on the moment of cue presentation Agren et al. [2012], Forcato et al. [2016]. We found that both RX and RCTX reminders reduced beta power in parieto-occipital regions, a pattern consistent with reactivation and updating of internal representations. Notably, sparse phase synchronization networks in the beta band were effective at distinguishing between the two types of reminders, indicating that the brain activates distinct functional architectures in response to retrieval reactivation to varying degrees. Crucially, the extent to which frontal nodes functioned as hubs, facilitating communication across the network, emerged as a significant factor: diminished frontal betweenness centrality correlated with poorer long-term behavioral outcomes. These findings underscore the importance of early network topology in influencing memory persistence, emphasizing that frontal regions play a vital role not only in retrieval and reactivation but also in the effective integration and dissemination of mnemonic information. Collectively, our results contribute to the growing body of evidence linking transient neural dynamics beyond the stimulus presentation to subsequent memory performance, thereby establishing a functional connection between momentary brain states and enduring behavioral outcomes.

## 4 Material and Methods

### 4.1 Participants

Thirty-three healthy undergraduate students from Pontificia Universidad Católica de Chile volunteered for the study (19 women). Their ages ranged from 18 to 35 years. All participants had to achieve at least 60% correct responses in the last four trials of the training session. Data from two subjects were excluded from analysis because they either did not complete the entire experiment or had excessively noisy EEG signals.

Before their participation, all subjects signed a written informed consent form. Both the protocol and the consent form were previously approved by the Ethics Committee of the Pontificia Universidad Católica de Chile, in accordance with the Declaration of Helsinki.

### 4.2 Experimental Design

We adapted the experiment used in Forcato et al. [2014]. The experiment consisted of three sessions separated by 24 hours, Figure 1A. On day 1, subjects learned a list of five pairs of nonsense syllables associated with a context (Training). The session on day 2 consisted of presenting reminders of the learned list and recording the neurophysiological data with an electroencephalogram (EEG). On day 3, we assessed the subjects’ memory (Testing).

The participants were divided into two groups based on the type of reminders they received on day 2.

Each trial consisted of presenting an image, playing classical music, and presenting the five pairs of syllables. In this sense, the list of syllables was associated with an enriched context (image + music).

#### 4.2.1 Training session

Each trial began with a black screen (3 seconds), followed by the context period, which included the presentation of an autumn forest image accompanied by Vivaldi’s music (5 seconds), and then the presentation of the list of syllables. The syllable period started with the presentation of a cue-syllable on the left-hand side of the screen and an empty box on the right side. Each cue-syllable was taken randomly from the list of five pairs. Subjects had 5 s to write the corresponding response syllable. After this, if no syllable was written, the correct one was shown for 5 s; if the response syllable was wrong, it was replaced by the correct one for 5 s; else, if the proper response was given, it stayed for 5 s longer. Immediately after that, another cue-syllable was shown, and the process was repeated until the list was over. Altogether, each trial lasted 58 s (3 s black screen, 5 s for context period, and 50 s for syllable presentation).

The training consisted of 10 trials. In each trial, we shuffle the presentation order of the pairs of syllables. In all the trials, participants were asked to write down the corresponding response syllable for each cue syllable presented. The list of five pairs of nonsense syllables was: **ITE**-*OBN*, **ASP**-*UOD*, **FLI**-*AIO*, **NEB**-*FOT*, **COS**-*GLE* (bold type: cue-syllable; italic type: response-syllable).

#### 4.2.2 Reminder session

Participants came to the same room used in the Training session. They sat in a chair in front of the computer, and we recorded their brain activity during a 4-minute resting period (*bl*_1_). Then, they received a reminder of the encoded material. Finally, we recorded the EEG signal during a 4-minute post-treatment resting period (*bl*_2_).

Participants were divided into two groups based on the reminder received. The RX group received the cue reminder (15 participants), while the RCTX group saw the context reminder (16 participants).

##### Cue reminder (RX group)

It consisted of presenting the context period (5 s, image + sound), followed by the presentation of the cue-syllable on the left-hand side of the monitor screen and the box on the right. After 5 seconds, the trial was interrupted, and an error message (‘ERROR 323’) appeared on the screen.

##### Context reminder (RCTX group)

It consisted of presenting the context period for 5 seconds (image + sound). Then, the trial was interrupted, and an error message (‘ERROR 323’) appeared on the screen.

#### 4.2.3 Testing session

The testing session consisted of 4 trials identical to the one presented during the Training session (each trial: 3 s black screen, 5 s for context period, and 50 s for syllable presentation). The testing session lasted 232 s.

### 4.3 Behavioral Data

We computed as a correct response every time a participant wrote down the correct response syllable per trial. To analyze long-term memory assessment, we calculated the mean of the correct responses throughout the four trials.

To compare the rate of correct responses of the two groups, we performed a Mann-Whitney test. To compare the variance between the groups, we used a Levene’s test.

To explore the relationship between long-term behavioral performance and neural markers, we divided each experimental group into two, based on the level of correct responses in the Testing session. We defined the low and high rate performers depending on whether their number of correct responses was below or above the median of all members in each experimental group (Figure SF1).

### 4.4 EEG data acquisition and preprocessing

EEG activity was recorded on a dedicated PC at a sampling frequency of 2048 Hz, with 64 Ag-AgCl active electrodes - mounted in an elastic cap according to the extended International 10–20 System, using the Biosemi Active-Two system (Biosemi, Amsterdam, the Netherlands, http://www.biosemi.com/products.htm). Two extra electrodes were placed at both mastoids. During the recording, impedances were kept below 20*k*Ω.

The preprocessing of the EEG recordings followed the pipeline used in Bavassi et al. [2017]. After acquisition, the data were imported into MATLAB using the EEGLAB toolbox Delorme and Makeig [2004]. First, they were digitally downsampled to 512 Hz using a fifth-order sinc filter to prevent aliasing, and referenced to the mean of the mastoid signal. The data were then bandpass filtered between 0.1 and 100 Hz using high-pass and low-pass filters, respectively. For this, we used the *eegfiltnew function* implemented in EEGLAB, which filters the signal forward and backward to achieve a zero-phase (flat frequency) response. Line noise, at 50 Hz, was removed by subtracting the evoked potential of the 50Hz signal from the original signal. To discard artifacts related to eye movements, we applied a global signal decomposition technique known as Independent Component Analysis (ICA), which is a source separation method that decomposes the EEG time series into a set of components aimed at identifying independent sources of variance in the data Delorme and Makeig [2004]. From the original 64 components, a subset of 1.9 *±* 0.9 was discarded. Finally, we applied a spherical channel interpolation at most 3 channels per subject after visual inspection.

### 4.5 Frequency decomposition and statistical analysis

After the data were preprocessed, we transformed all the neurophysiological time series into the time-frequency domain. For this analysis, we used the FieldTrip toolbox for MATLAB Oostenveld et al. [2011]. Specifically, we applied the Wavelet method, with 7 cycles, spanning the frequency range of 2 to 100 Hz. Power was calculated with moving steps of 0.5 s. Finally, the transformed data were averaged over the time dimension.

To study the modulation of the power oscillatory brain activity, we compared the activity before (*bl*_1_) and after (*bl*_2_) the presentation of the reminder, for each experimental group. A cluster-based statistical test with permutation correction was applied to address the multiple comparisons problem Maris and Oostenveld [2007]. This non-parametric test detects clusters of adjacent samples (channel, frequency) that show a statistically significant difference between *bl*_1_ and *bl*_2_. Clusters consisted of at least three neighboring channels and were considered significant within specific frequency bands. p-values were estimated using the Monte Carlo method, based on 500 random permutations (shuffling data between the two blocks), and the observed cluster statistics were compared against this distribution (the critical alpha level was set to 0.05).

Through this method, we identified frequency bands and electrode locations that simultaneously exhibited statistically significant activity.

We also performed a cluster-based permutation test to compare the two post-reminder power patterns (normalized by the pre-reminder resting), and we did not find any significant clusters.

### 4.6 Phase Synchronization Networks

As a measure of phase synchronization, we used the Phase Lag Index (*PLI*), which quantifies phase synchronization while avoiding spurious connectivity that may arise from volume conduction across the three-dimensional structure of the head Stam et al. [2007]. The *PLI* quantifies the contribution of coupling between two time series by detecting phase differences that are consistent over time but distinct from zero or *π*. To do this, the *PLI* computes the average sign of the instantaneous phase differences.

To build the synchronization matrices between electrode pairs (Figure 3A.) we applied a digital bandpass filter to the data within the frequency range identified by the non-parametric statistical analysis (25 − 40 Hz). Then, we downsampled the data to 128 Hz. Next, we applied a Hilbert Transform to each electrode’s time series, yielding a complex value for each channel *j* at time *t*, expressed as 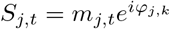 Demertzi et al. [2019]. *For each electrode pair j* and *k*, the *PLI* was computed as follows (Eq. 1) Chavez et al. [2010]:

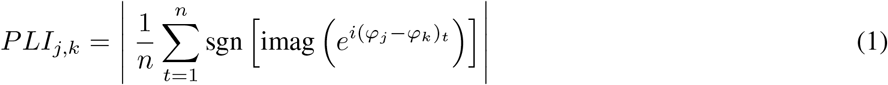

where *φ*_*j*_ and *φ*_*k*_ are the instantaneous phases of channels *j* and *k*, and sgn is the sign function, returning +1 if the argument is positive and − 1 if negative. This procedure yields a 64 *×* 64 symmetric matrix per subject and resting period 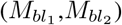.

To focus on the post-treatment synchronization pattern induced by the reminder, we normalized using the pre-treatment synchronization pattern per participant (Eq. 2):

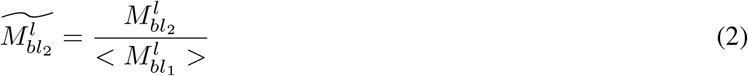

with 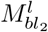 the synchronization matrix of the post-reminder resting period for subject *l* and 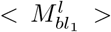 the mean synchronization of the pre-reminder for subject *l*.

Then, to compare between binary undirected networks, we thresholded the synchronization matrices for each participant to compare network’s attributes of graph with the same density (*ρ*), with density defined as the number of links over the total number of links 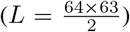. We fixed the number of edges, ranging from *ρ* = 1 to *ρ* = 10. Then, using Networkx (https://networkx.org/; a package for the creation, manipulation, and study of the structure, dynamics, and functions of complex networks), we calculated six network’s measures. We computed two node attributes: the node degree centrality (*k*_*i*_, number of neighbors of node *i*), and the betweenness centrality, which is the rate of shortest path (*d*_*u,v*_) that pass through node *i*. To simplify this local measures, we grouped the node centrality in four brain areas (Figure 3B): frontal right and left, and parietal right and left. Also, we computed global measures as the efficiency of the network (Eq. 3):

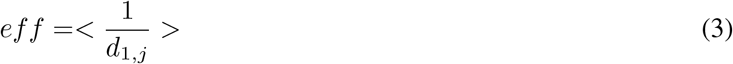

which is the mean of the inverse of the shortest path distance between the pairs (*d*_1,*j*_). Also, we computed the assortativity which is the Pearson’s Correlation between adjacent nodes (Barabási [2013]).Then, we evaluated the Giant component size, which quantifies the highest portion of connected nodes in the network (Barabási [2013]). We also calculated the global Clustering coefficient (Eq. 4):

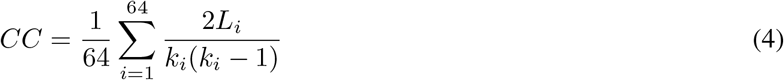

with *L*_*i*_ the number of connected neighbors of node *i* and *k*_*i*_ the node degree. To study the level of variability of each measure and compare them, we calculated the standard deviation over the mean of each measure 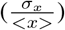.

To explore the existence of correlation between these network’s attribute, we computed the Pearson Correlation for each density. The correlation matrix in Figure 3D depicted the percentage of times two measures were significantly correlated (1: two magnitudes that were significantly correlated for all densities; 0: two measures that never were significantly correlated). We considered the correlation is significant if *p <* 0.05.

### 4.7 Classification Analysis and Feature Importance

For further analysis, we used Scikit-Learn, a library for machine learning in Python Pedregosa et al. [2011].

First, we applied a classification algorithm to distinguish between the networks and the RX and RCTX reminder groups based on the local and global attributes that presented a higher level of variability and low correlation between them (efficiency, clustering, and the four betweenness centralities). Considering the low amount of data, we only had 31 networks per density, we performed a Random Forest Classifier, which is a robust algorithm that does not need data scaling, and works well with a small amount of data. The Random Forest classifier consisted of 50 shallow trees (maximum depth = 2), with only two features considered at each split, to minimize overfitting. To assess the classifier’s performance, we employed Leave-One-Out Cross-Validation (LOOCV). We computed two evaluation metrics: macro-averaged F1 score and overall accuracy. Both training and test scores were recorded across all LOOCV folds. Final performance (Figure 4A) was reported as the average and standard error of the cross-validated scores across all subjects. This allowed us to estimate the model’s generalization performance while ensuring robustness to data imbalance.

Then, to capture the most relevant network’s measure in distinguishing between both groups, in the densities where the previous classifier was above chance, we applied a Random Forest-based feature selection. We evaluated feature importance using a permutation-based approach. The data were randomly split into a training set (70%) and a test set (30%), and the Random Forest model was fitted to the training data. Feature importance was then assessed using the permutation importance method with 50 random shuffles per feature, computed on the held-out test set. For each feature, we stored both the mean importance and its standard error. In parallel, for each density, we performed automatic feature selection using a model-based method. The trained Random Forest model provides internal estimates of feature importance based on the Gini impurity reduction associated with each split. We used scikit-learn’s *SelectFromModel* function, which selects features whose importance scores exceed a predefined threshold (by default, the mean importance across all features). The selected features, frontal right and left betweenness, presented scores above the threshold in 95% of the densities.

Finally, we aimed to identify low vs. high performance during the Testing session based on bilateral frontal centrality. So, for each group, we applied a generalized linear model (GLM) with a binomial distribution and logit link function was fitted to assess the contribution of left and right frontal centrality, and the connectivity density to predict the binary outcome variable (Low/High). The analysis was conducted in Python using the statsmodels package. A constant term was included in the model to estimate the intercept, and parameters were optimized using the Iteratively Reweighted Least Squares (IRLS) method. To evaluate the discriminative capacity of the models, we calculated the area under the receiver operating characteristic (ROC) curve for each one. Then, we analyzed whether the observed model performance was above chance level; therefore, we generated a null distribution of AUC values using a randomization procedure. We randomly permuted 1000 times the outcome variable (Low/High), and for each iteration, a logistic GLM with the same predictors was refitted using the randomized labels.

## Supporting information

https://drive.google.com/file/d/15qPZuylJhhEBgtnOJ6UnooIHX9j-XoMl/view?usp=sharing

