## Supplementary material for "The hidden phase of memory: EEG signatures of reactivation during post-retrieval rest": https://drive.google.com/file/d/15qPZuylJhhEBgtnOJ6UnooIHX9j-XoMl/view?usp=sharing

A PREPRINT

**Luz Bavassi**
Instituto de Fisiología, Biología Molecular y Neurociencias (IFIBYNE),
CONICET, Buenos Aires, Argentina
Departamento de Física, Universidad de Buenos Aires,
Buenos Aires, Argentina


**Germán Campos- Arteaga**
Departamento de Ciencias Jurídicas y Sociales, Facultad de Ciencias Jurídicas y Sociales,
Universidad Tecnológica Metropolitana, Santiago, Chile.


**Ismael Palacios-García**
Centro de Estudios en Neurociencia Humana y Neuropsicología,
Facultad de Psicología, Universidad Diego Portales, Santiago, Chile


**Mario Villena-Gonzalez**
Departamento de Ciencias Jurídicas y Sociales, Facultad de Ciencias Jurídicas y Sociales,
Universidad Tecnológica Metropolitana, Santiago, Chile.


**Libertad Campassi**
Departamento de Física, Universidad de Buenos Aires,
Buenos Aires, Argentina


**Emiliano Marachlian**
Instituto de Fisiología, Biología Molecular y Neurociencias (IFIBYNE), CONICET, Buenos Aires, Argentina
Departamento de Física, Universidad de Buenos Aires,
Buenos Aires, Argentina


**Cecilia Forcato**
Laboratorio de Sueño y Memoria, Instituto Tecnológico de Buenos Aires (ITBA)
Buenos Aires, Argentina


**Eugenio Rodriguez Balboa**
Pontificia. Universidad Católica de Chile, Laboratorio de Neurodinámica Básica y Aplicada,
Escuela de Psicología, Santiago, Chile.


**Eugenia Pedreira**
Instituto de Fisiología, Biología Molecular y Neurociencias (IFIBYNE),
CONICET, Buenos Aires, Argentina


### ABSTRACT

Memory retrieval reactivates previously encoded representations, allowing for their modification and strengthening. However, the neural processes that follow reactivation and contribute to long-term retention remain poorly understood. Here, we examined the post-retrieval resting period to identify neural markers of memory reactivation and their relation to subsequent memory performance. Participants learned pairs of nonsense syllables in multisensory contexts and, on the following day, received either a cue-syllable reminder (context + cue syllable; RX) or a context-only reminder (RCTX) while EEG activity was recorded before and after the reminder presentation. Both reminders elicited significant reductions in beta power (25–40 Hz) during the post-reminder rest, consistent with memory reactivation. The magnitude of beta decrement correlated with better long-term performance. Graph-theoretical analyses of phase synchronization networks in the beta band revealed that the RCTX reminder produced higher bilateral frontal betweenness centrality, suggesting greater engagement of frontal regions in mediating global information flow when retrieving only contextual cues. Moreover, frontal centrality and network density were predictive of subsequent memory accuracy. These findings demonstrate that memory reactivation extends beyond cue presentation into post-retrieval rest, leaving identifiable oscillatory and topological signatures that influence memory persistence. Our results underscore the crucial role of frontal network dynamics and beta-band activity in facilitating long-term memory.


***Keywords*** Resting State · Memory reactivation · Beta power reduction · Subsequent memory accuracy · Phase synchronization network

### 1 Low vs. High performers in the Testing Session

We split participants into two groups depending on their performance accuracy in the Testing Session. Participants whose accuracy was above the median accuracy of the whole group were assigned as High performers, while those below or equal to the median were classified as Low performers. Figure 1 depicts the distribution of Low performers (blue) and High performers in both experimental groups.

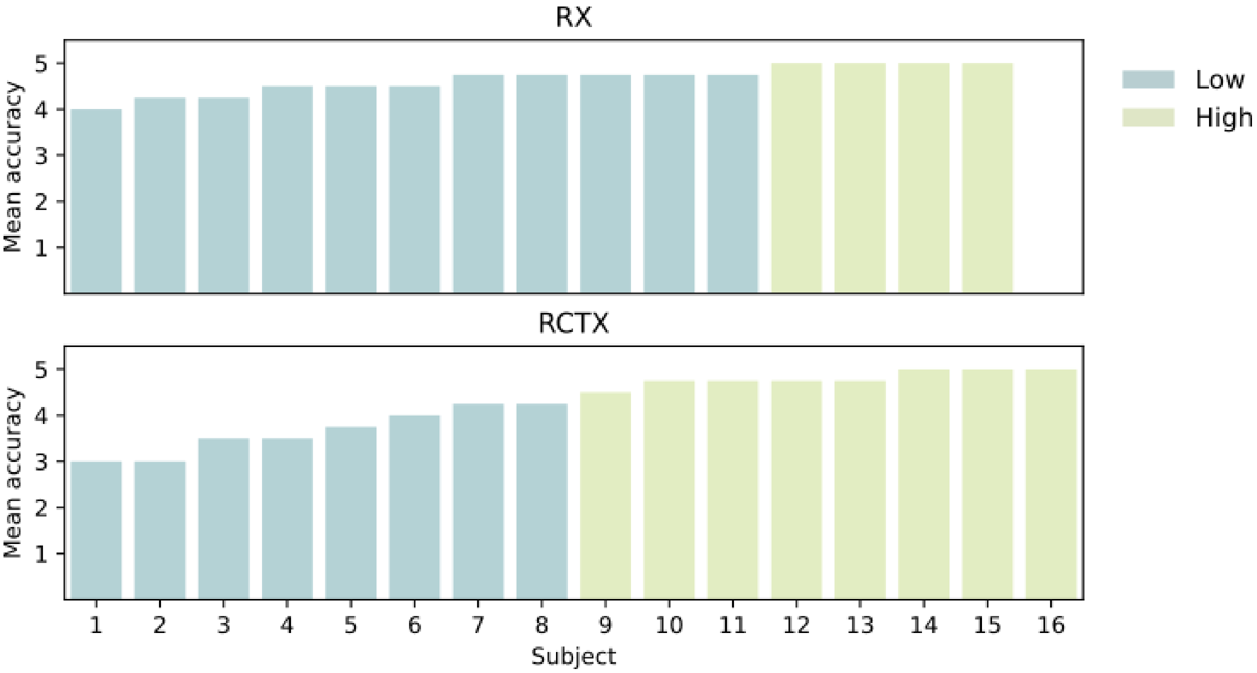


Figure 1: **Low and High performers.** Participants were split into two subgroups depending on their accuracy in the Testing Session. For the RX reminder group (top panel), the median was 4.75, while for the RCTX group (bottom panel), the median was 4.38 (Low: blue; High: light green).

### 2 An alpha power significant cluster in the RX group

When we compared the resting period post-retrieval ($bl_2$) with the pre-retrieval period ($bl_1$) using a permutation cluster-based analysis in the RX reminder group, a positive cluster emerged around alpha power ($7 - 12$ Hz). This cluster is distributed throughout the brain, predominantly on the right side, spanning from the frontal to the occipital regions (Figure 2A, $p = 0.05$, $T = 293.8$, cluster-based permutation test). Then, we examined whether this difference was related to the subsequent performance (Figure 2B). There were no differences between the Low and High performers ($U = 24$, $p = 0.85$; Mann–Whitney test).

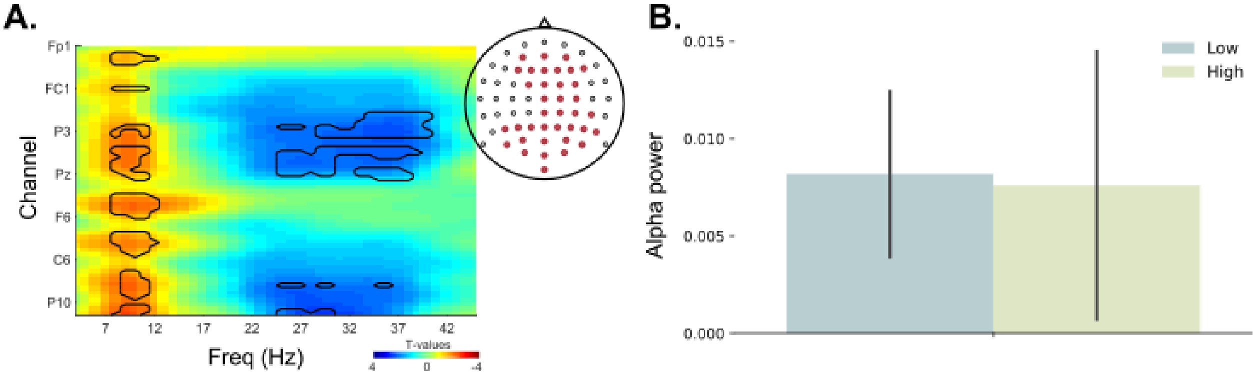


Figure 2: **Spectral power analysis for RX. A.** T- maps of spectral power changes between $bl_2$ and $bl_1$ in the RX group. The thick black line depicts the significant cluster ($p < .05$, cluster-based permutation test). On the left, the location of EEG-channels for the alpha cluster. **B.** The alpha power ($bl_2 - bl_1$) of the significant cluster for High and Low performance in the Testing Session. (Low: blue; High: light green).

### 3 Distance between phase synchronization networks.

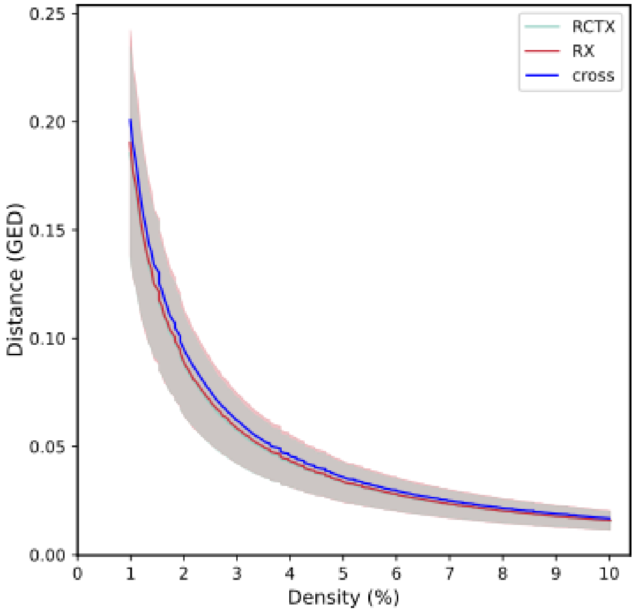


Figure 3: **Distance curves.** Graph Edit distance within each experimental group (RX: red; RCTX: green) and between groups (blue). The three curves are indistinguishable along the density axis.

Once we had built the phase synchrony networks for each participant at each connectivity density, we compared them to assess their similarity. We computed the distance within each group and the distance between the two groups. For this, we used the Graph Edit Distance (GED) Emmert-Streib et al. [2016], which compares two graphs in an error-tolerant manner, computing the series of edit operations that transform a graph $\mathcal{G}_1$ into $\mathcal{G}_2$ by producing minimal transformation costs as the optimal inexact match. Finally, the graph edit distance of two given graphs is the minimum cost associated with a series of edit operations.

We calculated the distance between every pair of networks for each density (Figure 3), and it was not possible to assign them to a specific group because the three curves were almost identical.
